# Optimization of Phosphate Limited Autoinduction Broth for 2-Stage Heterologous Protein Expression in *E. coli*

**DOI:** 10.1101/2021.06.08.447502

**Authors:** Romel Menacho-Melgar, Jennifer N. Hennigan, Michael D. Lynch

**Affiliations:** Department of Biomedical Engineering, Duke University; Department of Chemistry, Duke University

**Keywords:** Protein Expression, Autoinduction media, Phosphate induction

## Abstract

Autoinducible, 2-stage protein expression leveraging phosphate inducible promoters has been recently shown to enable not only high protein titers but also consistent performance across scales from screening systems (microtiter plates) to instrumented bioreactors. However, to date small scale production using microtiter plates and shake flasks rely on a complex autoinduction broth (AB) that requires making numerous media components, not all amenable to autoclaving. In this report, we develop a simpler media formulation (AB-2) with just a few autoclavable components. We show that AB-2 is robust to small changes in its composition and performs equally, if not better, than AB across different scales. AB-2 will facilitate adoption of phosphate limited 2-stage protein expression protocols.

## Introduction

Heterologous protein expression is a ubiquitous workflow used in numerous labs performing biological research. (1) Additionally, *E. coli* is still a key workhorse host for protein expression and purification. (2, 3) We have recently reported improved strains and plasmids for tightly controlled expression upon phosphate limitation in 2-stage cultures (Figure 1). (4, 5) This method requires placing the gene for the protein of interest under the control of a phosphate inducible promoter which becomes activated under low-phosphate conditions through the PhoRB two-component signal transduction system.(6, 7) Further, we have evaluated a comprehensive set of such promoters and characterized a few to be robust to media composition and scalable from microtiter plates to instrumented bioreactors in commercially relevant minimal media (8) In this 2-stage expression method, protein expression is induced upon entry into stationary phase when phosphate becomes depleted, thereby decoupling cell growth from protein production. Decoupling growth from protein production lessens the metabolic burden of heterologous protein expression on growth, facilitating the adoption of standardized protocols across proteins of interest and scales.(9) Our 2-stage protocol further eliminates the need to optimize induction conditions while maintaining high protein titers; which improves workflows when compared to routinely-used growth-associated expression protocols.(10, 11)

**Figure 1:**
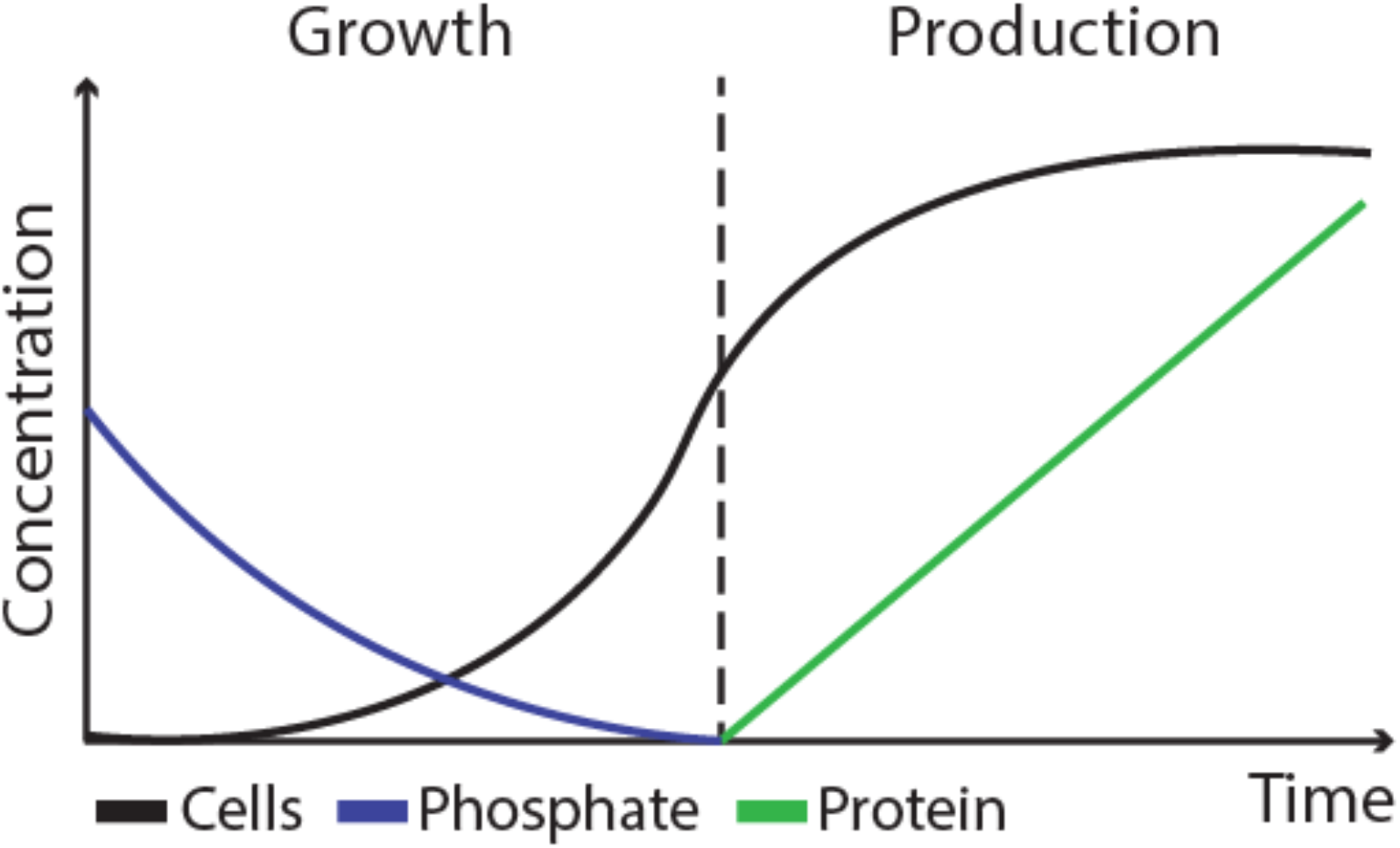
2-stage autoinducible heterologous expression upon low phosphate induction. Cells (black line) are grown during the growth phase until phosphate in the media (blue line) becomes depleted. Phosphate depletion activates low phosphate promoters which are used to drive expression of heterologous protein (green line) during the production phase.

These strains and plasmids offer robust expression, which is readily scalable from micro-titer plates to instrumented bioreactors.(4, 8) Additionally, several of these strains enable rapid autolysis and auto DNA/RNA hydrolysis simplifying protein purification.(12) While expression in instrumented fermentations uses defined minimal mineral salts media, small scale production of proteins using these strains and protocols relies on an optimized phosphate limited Autoinduction Broth (AB).(4) The initial formulations of this media are quite complex and require the preparation of 11 components from 17 ingredients, several of which cannot be autoclaved. This complexity limits the ease of adopting these new protocols. To address this limitation, in this study, we report the development of a simpler autoinduction broth (AB-2), with far fewer components and a simpler preparation, which allows for equivalent protein expression levels.

## Materials & Methods

### Chemicals and Media Components

MOPS and Bis-Tris were purchased from GoldBio (St Louis, MO). Yeast extract and Casamino acids were purchased from BioBasic (Amherst, NY). Ammonium sulfate anhydrous and glucose were purchased from VWR (Radnor, PA). All other reagents were purchased from Sigma-Aldrich (St. Louis, MO). Stock solutions used in this study are shown in Supplemental Table 1. All components were autoclaved except for CaSO_4_, MgSO_4_, thiamine-HCl, Trace Metals and ferric sulfate which were filter sterilized. Glucose, MOPS and Bis-Tris were filter sterilized unless otherwise stated.

### Plate and Flask part numbers

250 ml Erlenmeyer flasks were purchased from VWR (Cat#89095-270), 96 well microtiter plates were purchased from Genesee Scientific (San Diego, CA; Cat#25-104) and 384 well plates were purchased from VWR (Cat#10814-224).

### Strains and Plasmids

Strain DLF_R002 and plasmid pHCKan-yibDp-GFPuv (Addgene #127078) were constructed as described previously. (4)

### Media Preparation

Autoinduction broth (AB) was prepared as described previously.(4) AB media variations used in initial experiments to simplify media composition were prepared in the same manner as AB, but dropping the indicated media components. To prepare media used in experiments to evaluate the effect of autoclaving glucose and buffers, all components were mixed from stock solutions and autoclaved except when noted, in which case the component was filter-sterilized and added to the cooled-down autoclaved media mixture. To make AB-2 medium, 6.2 g of yeast extract, 3.5 g of casamino acids, 5.4 g of ammonium sulfate anhydrous and 41.8 g of Bis-Tris were mixed and water added to a volume of 907 mL. 3 ml of 12 M HCl was then added to adjust pH. A second solution of 500 g/L glucose was made and autoclaved. After both solutions were cooled down, 90 ml of the 500 g/L glucose solution was added to the first mixture to make AB-2. For AB-2 robustness studies, media were made according to the AB 2.0 formulation, but leaving the component being varied out. Then the missing component was added beginning with 75% and ending at 125% at 5% increments to the media and aliquoted for use at each increment for testing.

### Micro-fermentations

Micro-fermentations were performed as described previously.(4) Briefly, a 5 ml of LB culture was started from a frozen stock of DLF_R002 with pHCKan-yibDp-GFPuv and grown overnight at 37 °C and 150 rpm. The overnight culture was used to inoculate at 1% v/v 100 μl and 15 μl of media per well in 96 well and 384 well plates, respectively. AB-2 was supplemented with 0.05% of polypropylene glycol (2000 Da MW) for 384 well plate micro-fermentations. Micro-fermentations were incubated at 37 °C and 300 rpm for 24 hours. Sandwich covers were used to prevent evaporation and were purchased from EnzyScreen (Haarlem, The Netherlands; 96 well plate cover Cat#CR1596 and 384 well plate cover Cat#CR1384).

### BioLector Studies

Biolector studies were conducted as described previously.(4) Briefly, an overnight LB culture of DLF_R002 with pHCKan-yibDp-GFPuv was normalized to 25 OD (600 nm) and used to inoculate at 1% v/v 792 μl of AB or AB-2 in a Flower-Plate (Cat no MTP-48-B, m2p-labs). Growth and fluorescence measurements were obtained using a BioLector (m2p-labs, Baesweiler, Germany) over 36 hours using a 20 biomass gain, 20 GFP gain (488 nm excitation and 620 nm emission), 1300 rpm, 37 °C and 85% humidity.

### Shake Flask Production

An overnight LB culture of DLF_R002 with pHCKan-yibDp-GFPuv was used to inoculate 20 ml of AB or AB-2 in vented baffled Erlenmeyer flasks and incubated at 37 °C and 150 rpm for 24 hours.

### OD and GFP measurements

OD600 and GFP measurements were made using a Tecan Infinite 200 plate reader and 200 ul black-walled plates purchased from Fisher Scientific (Hampton, NH; Cat#07-000-136) at a 40-fold dilution. OD was read at 600 nm and GFP was measured using excitation at 412 nm and emission at 530 with filters purchased from Omega Optical (Brattleboro, VT; Cat#3019445, #3024970 and #3032166, respectively).

## Results & Discussion

Our original autoinduction broth (AB) was derived from a minimal mineral salts media (FGM10) used in instrumented bioreactors. (4) As a result, AB retained the components used in minimal media including numerous salts and trace metals. AB was modified from FGM10 to include complex nutrient sources (yeast extract and casamino acids) that were added to reduce the lag phase in smaller batch cultures in microtiter plates and shake flasks. (Figure 2a) As these complex nutrient sources also are expected to supply many salts and trace metals, we first sought to evaluate the impact of dropping out these individual components (alone and in combination) on growth and protein expression. Toward this goal, we measured endpoint biomass (OD_600nm_) and GFP levels (GFPuv) in modified AB missing several components as illustrated in Figure 2a. In these experiments, GFP was expressed from the low phosphate inducible *yibDp* promoter from pHCKan-yibDp-GFPuv (Addgene 127078), using *E. coli* strain DLF_R002. DLF_R002 has deletions for *ackA-pta*, *pflB*, *adhE*, *ldhA* and *poxB* which reduce production of mixed acid fermentation products from overflow metabolites, including acetate which can lower growth and protein production. (4, 13, 14) Additionally, DLF_R002 has deletions of *iclR* and *arcA* which result in increased biomass yield and reduced overflow metabolism.(4, 15) Together, the modifications in DLF_R002 completely eliminate organic acid byproducts, including acetate, even under excess glucose fermentation conditions.(4) Cultivations using DLF_002 with pHCKan-yibDp-GFPuv were performed in 96 well microtiter plates. As seen in Figure 2a, a formulation consisting of only glucose, buffer, ammonium sulfate, yeast extract and casamino acids resulted in comparable expression levels with the original AB, greatly simplifying the media formulation.

**Figure 2:**
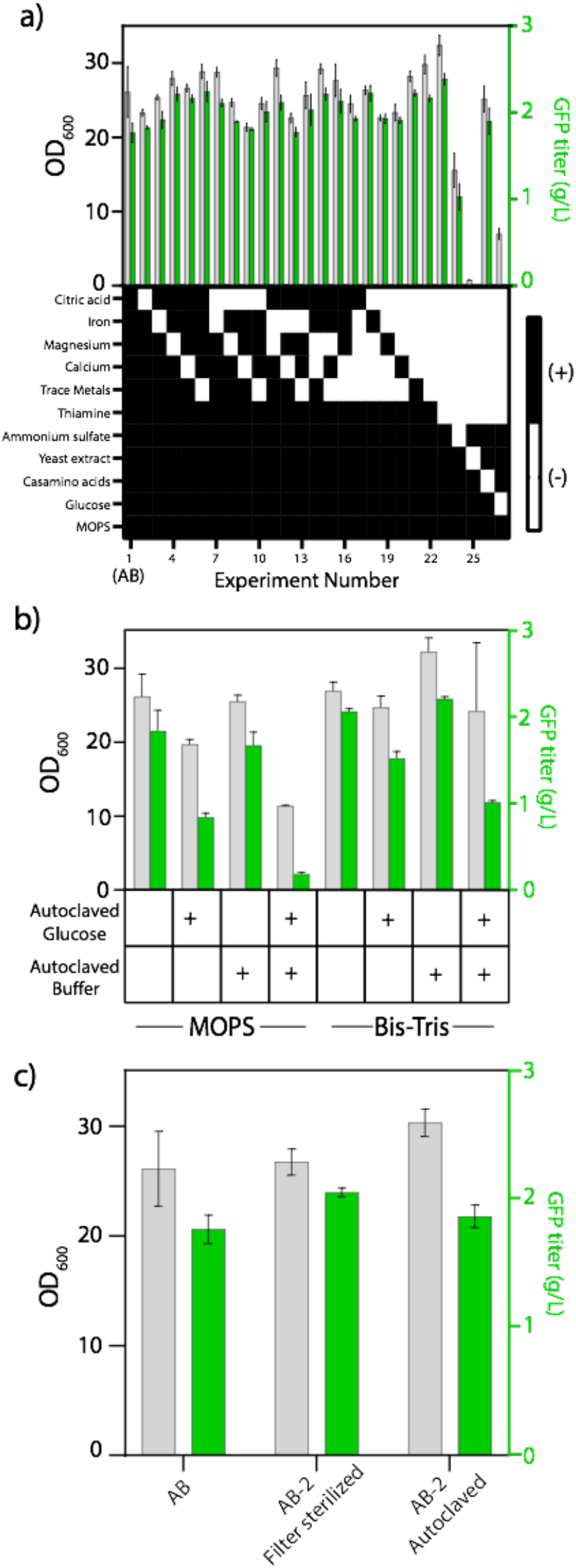
Autoinduction media optimization. For all panels, green bars correspond to average GFP titer (g/L) and grey bars indicate average biomass (OD_600_) resulting from triplicate experiments. (a) Autoinduction media variations with respect to AB (first data point) resulting from dropping various components from the AB recipe. The media composition for each medium is indicated in the heat map immediately below the data bars showing black if the component is present or white if it is not. (b) Effect of buffer selection and autoclaving glucose and/or buffer in AB-2 on biomass and GFP production. (c) AB comparison with AB-2 prepared by mixing components and filter sterilizing (‘AB-2 Filter sterilized’) or by weighing media components and sterilization by autoclaving where glucose was autoclaved separately from other components (‘AB-2 Autoclaved’).

Another limitation to the original AB is the use of 3-morpholinopropane-1-sulfonic acid (MOPS) as a buffer. While MOPS buffers in the appropriate range (pH 6.5-7.9), autoclaving or prolonged storage at room temperature of sulfonic acid buffers is not recommended because they produce unknown yellow degradation products.(16) To overcome this issue, we evaluated the use of an alternative buffer that is completely stable during sterilization namely Bis-Tris.(17) As can be seen in Figure 2b, replacing the buffer with Bis-Tris, led to equivalent growth and expression, while enabling the buffer to be mixed with the ammonium sulfate, yeast extract and casamino acids prior to sterilization by autoclaving.

By utilizing Bis-Tris as a buffer, we consolidated the AB media to only two components, i) a concentrated glucose solution and ii) a buffer, ammonium sulfate, yeast extract and casamino acid mixture, both of which can be sterilized by autoclaving. Next, we investigated the ability to mix the glucose with the other components prior to sterilization. While autoclaving glucose with amino acids present in complex nutrient sources can lead to Maillard reactions products, (18) which have been shown to inhibit growth in some applications, (18, 19) the toxicity of these products can be strain dependent.(19) Thus, we evaluated single mixture formulations, results of which are also given in Figure 2b. Unfortunately, as with many other media, glucose needs to be autoclaved separately from the other components for optimal growth and autoinduction of protein expression. We additionally showed AB-2 made from powder components and autoclaved performs similarly to AB or AB-2 made from pre-sterilized stock solutions as shown in Figure 2c. We settled on a final AB-2 formulation (Table 1) and compared the time course of cellular growth and GFPuv auto-induction in AB vs AB-2 (Figure 3).While some differences in the growth curves are observed during mid-stationary phase, the growth and protein production levels are equivalent or improved in AB-2 over AB.

**Table 1:** AB-2 Media Formulation.

|  | Glucose Solution 1<br>(per L) | Solution 2 (per L) |
| --- | --- | --- |
| Glucose | 500g | -- |
| Bis-Tris | -- | 41.8 g |
| Ammonium Sulfate anhydrous | -- | 5.4 g |
| Yeast Extract | -- | 6.2 g |
| Casamino Acids | -- | 3.5 g |
| 12 M HCl | -- | 3 mL |
| Deionized Water | QS to 1L | QS to 910 mL |
| Sterilize | Autoclaving | Autoclaving |
| For 1 L of AB-2, add 90 mL of sterilized Glucose Solution 1 to 910 mL of sterilized Solution 2. |  |  |

**Figure 3.**
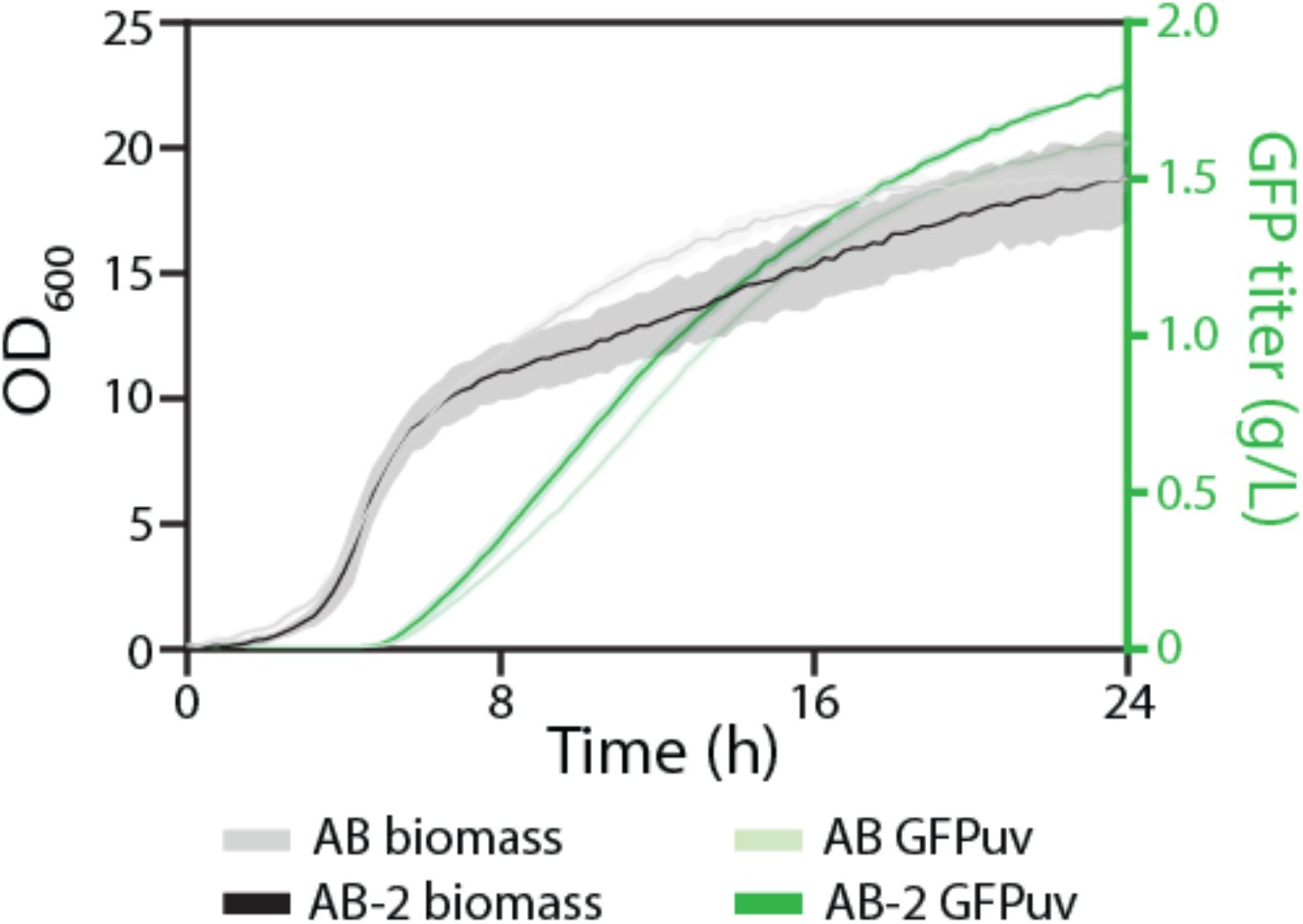
Growth and GFP production curves for AB and AB-2 media. Black lines correspond to average growth curves and green lines correspond to average GFP titer resulting from (n=4) experiments. Lighter lines correspond to AB medium cell cultures and dark lines correspond to AB-2 medium cell cultures. Shaded areas correspond to the standard deviation.

With the success in simplifying the formulation and preparation of AB-2, we sought to evaluate how sensitive this formulation was to small changes in the levels of various components. Toward this goal we changed the concentration of each individual component to its 75% (or −25%) up to its 125% (or +25%), at 5% increments. Results of these studies are given in Figure 4. Briefly, growth and auto-induction in AB-2 is quite robust to errors in preparation. We next performed a head to head comparison of AB and AB-2 at multiple production scales, including 384 well plates, 96 well plates, 250 mL shaker flasks (20mL fill volume). Again these studies used strain DLF_R002, expressing GFPuv as described above. Results are given in Figure 5. In these studies, we demonstrated that AB-2 performs as well as AB in terms for growth and protein expression, across scales.

**Figure 4.**
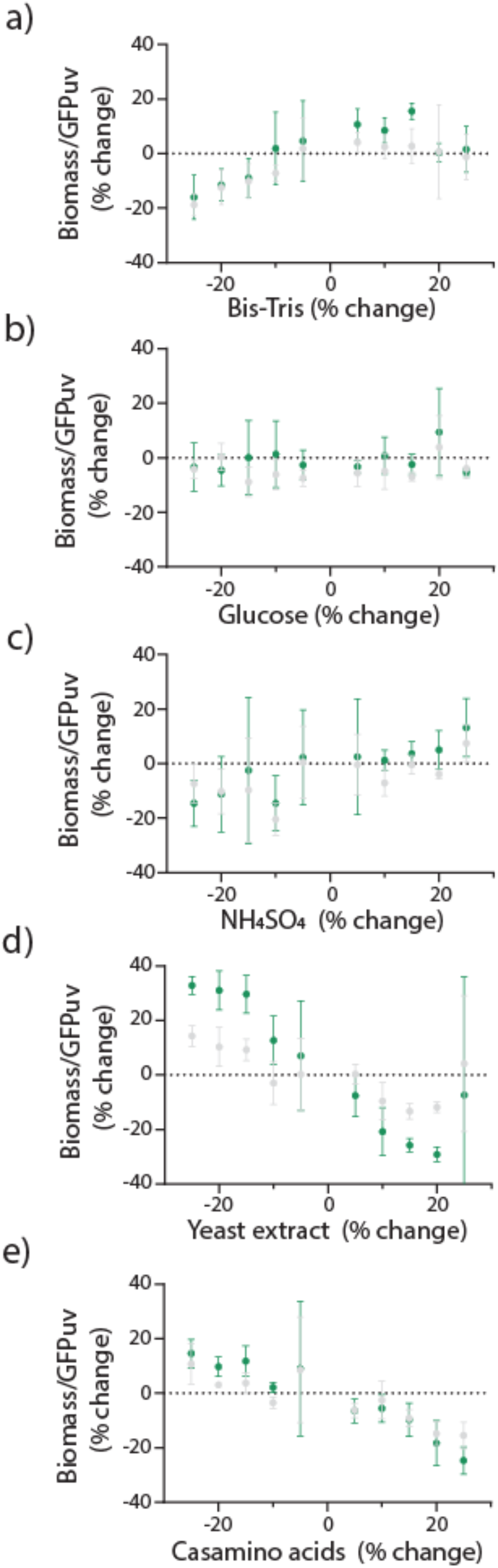
Effect of changing the concentrations of one component at a time in AB-2 from −25% to +25%, at 5% intervals, on biomass (gray circles) and GFPuv (green circles). Effect of changes in (a) Bis-Tris, (b) glucose, (c) ammonium sulfate, (d) yeast extract and (e) casamino acids. All experiments were done in triplicate.

**Figure 5:**
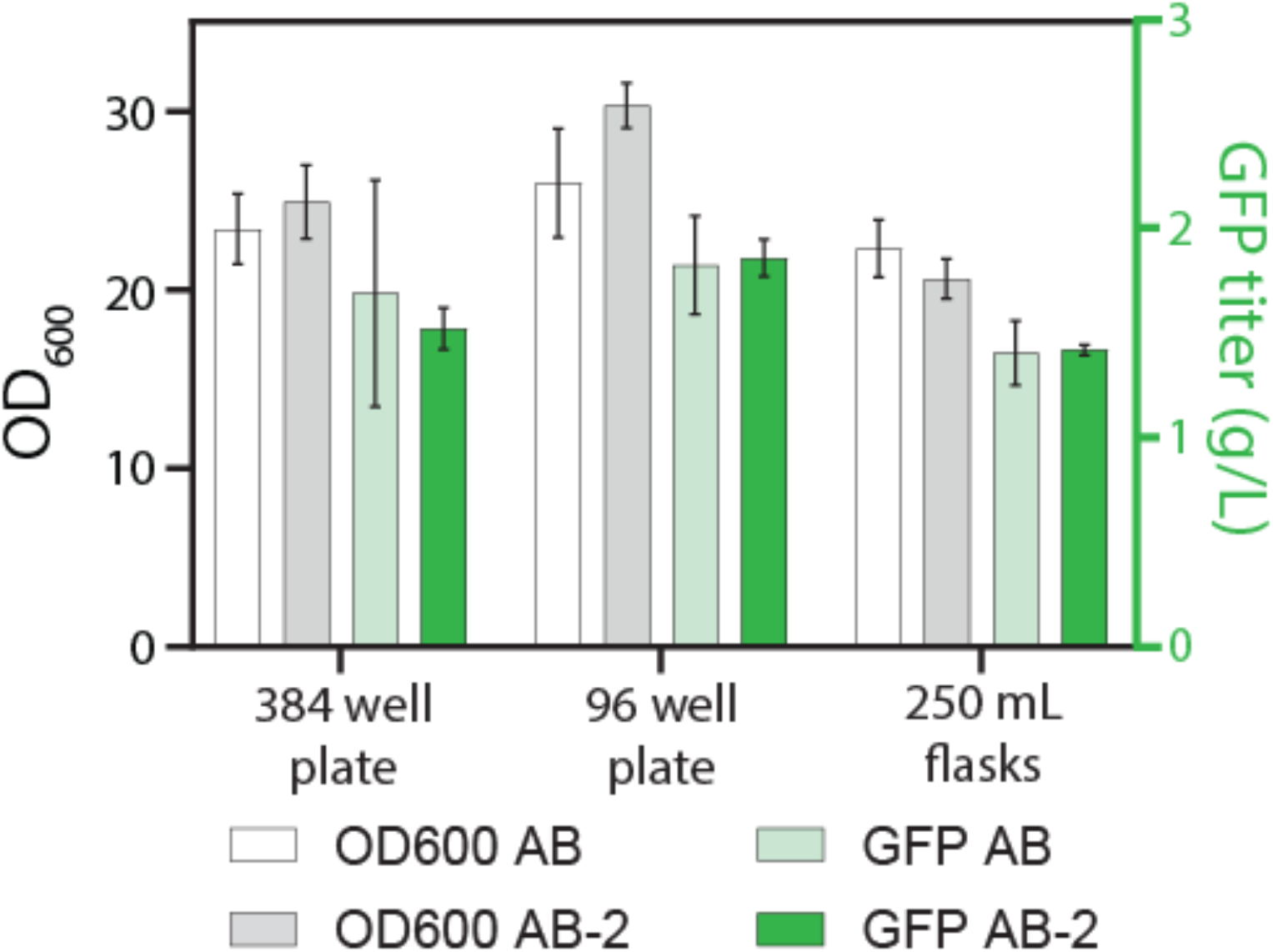
AB-2 performance across scales compared to AB after 24 hours. White and grey bars correspond to biomass levels in AB and AB-2, respectively. Light and dark green bars correspond to GFP titer in AB and AB-2, respectively. 15 μl, 100 μl and 20 ml fill volumes were used for 384 well plates, 96 well plates and 250 ml Erlenmeyer flasks, respectively. Results shown are averages from triplicate experiments.

The new AB-2 formulation is simpler and less expensive than the original AB while offering near equivalent (if not improved) growth and protein expression. Importantly, the new formulation is extremely simple to prepare. Key components can be mixed as dry powders, and after the addition of water, autoclaved as only two components. This new media will simplify the adoption of phosphate limited autoinduction for 2-stage heterologous protein expression in *E. coli.*

## Acknowledgements

We would like to acknowledge the following support: North Carolina Biotechnology Center 2018-BIG-6503. R. Menacho-Melgar was supported in part by the Entrepreneurial Post-Doctoral fellowship from the Department of Biomedical Engineering, Duke University.

## Author contributions

R. Menacho-Melgar designed, executed and analyzed experiments. J.N.Hennigan performed microfermention experiments. M.D. Lynch designed and analyzed experiments. All authors wrote, revised and edited the manuscript.

