## Supplementary Materials for "Optimization of Phosphate Limited Autoinduction Broth for 2-Stage Heterologous Protein Expression in *E. coli*"

**Supplemental Materials**

| **Supplemental Table 1: Media stock solutions used in this study** | | |
|---|---|---|
| **Component** | **Composition per 1 L** | **Stock concentration** |
| Ammonium sulfate anhydrous | -396.42 g of ammonium sulfate | 3 M |
| Citric acid | -100 of citric acid | 100 g/L |
| $FeSO_4 \cdot 7H_2O$ | -27.8 g of $FeSO_4 \cdot 7H_2O$<br>-2.5 ml of concentrated $H_2SO_4$ | 100 mM |
| $MgSO_4$ | -492.96 of $MgSO_4 \cdot 2H_2O$ | 2 M |
| $CaSO_4$ | -1.72 g of $CaSO_4 \cdot 2H_2O$ | 10 mM |
| Glucose | -500 g of glucose | 500 g/L |
| MOPS | -209.3 g of MOPS | 1 M |
| Bis-Tris | -209.2 g of Bis-Tris | 1 M |
| Thiamine-HCl | -50 g of thiamine-HCl | 50 g/L |
| Yeast Extract | -100 of yeast extract | 100 g/L |
| Casamino Acids | -100 of casamino acids | 100 g/L |
| Trace Metals | -10 mL of concentrated $H_2SO_4$<br>-0.6 g $CoSO_4 \cdot 7H_2O$<br>-0.5 g $CuSO_4 \cdot 5H_2O$<br>-0.6 g $ZnSO_4 \cdot 7H_2O$<br>-0.2 g $Na_2MoO_4 \cdot 2H_2O$<br>-0.1 g $H_3BO_3$<br>-0.3 g $MnSO_4 \cdot H_2O$ | 500 X |
